# Altered chromatin localization of hybrid lethality proteins in *Drosophila*

**DOI:** 10.1101/438432

**Authors:** J.C. Cooper, A. Lukacs, S. Reich, T. Schauer, A. Imhof, N. Phadnis

## Abstract

Understanding hybrid incompatibilities is a fundamental pursuit in evolutionary genetics. In crosses between *Drosophila melanogaster* females and *Drosophila simulans* males, the interaction of at least three genes is necessary for hybrid male lethality: *Hmr ^mel^*, *Lhr ^sim^*, and *gfzf ^sim^*. All three hybrid incompatibility genes are chromatin associated factors. While HMR and LHR physically bind each other and function together in a single complex, the connection between either of these proteins and *gfzf* remains mysterious. Here, we investigate the allele specific chromatin binding patterns of *gfzf*. First, our cytological analyses show that there is little difference in protein localization of GFZF between the two species except at telomeric sequences. In particular, GFZF binds the telomeric retrotransposon repeat arrays, and the differential binding of GFZF at telomeres reflects the rapid changes in sequence composition at telomeres between *D. melanogaster* and *D. simulans*. Second, we investigate the patterns of GFZF and HMR co-localization and find that the two proteins do not normally co-localize in *D. melanogaster*. However, in inter-species hybrids, HMR shows extensive mis-localization to GFZF sites, and this altered localization requires the presence of *gfzf ^sim^*. Third, we find by ChIP-Seq that over-expression of HMR and LHR within species is sufficient to cause HMR to mis-localize to GFZF binding sites, indicating that HMR has a natural low affinity for GFZF sites. Together, these studies provide the first insights into the different properties of *gfzf* between *D. melanogaster* and *D. simulans* as well as a molecular interaction between *gfzf* and *Hmr* in the form of altered protein localization.

## Introduction

Speciation, the process by which one species splits into two, involves the evolution of reproductive isolating barriers between previously interbreeding populations (Dobzhansky 1937; Coyne and Orr 2004). Intrinsic postzygotic barriers, such as the sterility or inviability of hybrids, are caused by deleterious genetic interactions known as hybrid incompatibilities. Understanding the evolutionary forces that generate such barriers between species requires understanding the properties of incompatibility genes – namely, what are their native functions within species, and how do they interact with each other? Despite decades of studies, a system of species with detailed molecular understanding of their hybrid incompatibilities remains elusive.

One of the longest studied inter-species hybridizations is between the fruit fly *Drosophila melanogaster* and its closest sister species, *Drosophila simulans*. When *D. melanogaster* females are crossed to *D. simulans* males, the hybrid F1 males die during larval development (Sturtevant 1919). These F1 hybrid males lack imaginal disc tissues, and do not undergo sufficient growth to trigger pupation; hybrid F1 females live to adulthood but are sterile (Sánchez and Dübendorfer 1983). Three genes are essential for the inviability of these hybrid F1 males: *Hybrid male rescue* from *D. melanogaster* (*Hmr ^mel^*) (Hutter and Ashburner 1987; Barbash *et al.* 2003), *Lethal hybrid rescue* from *D. simulans* (*Lhr ^sim^*) (Watanabe 1979; Brideau *et al.* 2006), and GST-containing FLYWCH zinc-finger protein from *D. simulans* (*gfzf ^sim^*) (Phadnis *et al.* 2015) (Figure 1). These three alleles interact genetically to form a single hybrid incompatibility, and all three incompatible alleles must be simultaneously present for hybrid F1 male lethality. Loss-of-function or reduced expression of any individual incompatible allele is sufficient to rescue the viability of hybrid F1 males. *Hmr ^mel^* is lethal to hybrid males while *Hmr ^sim^* does not cause hybrid male inviability. Conversely, *Lhr ^sim^* and *gfzf ^sim^* are incompatible alleles that cause hybrid F1 male lethality, whereas the corresponding alleles of *Lhr ^mel^* and *gfzf ^mel^* do not contribute to hybrid F1 male lethality. The *D. melanogaster-D. simulans* hybridization thus represents one of the best understood systems in terms of the identities of the genes involved in hybrid lethality (Barbash 2010).

**Figure 1.**
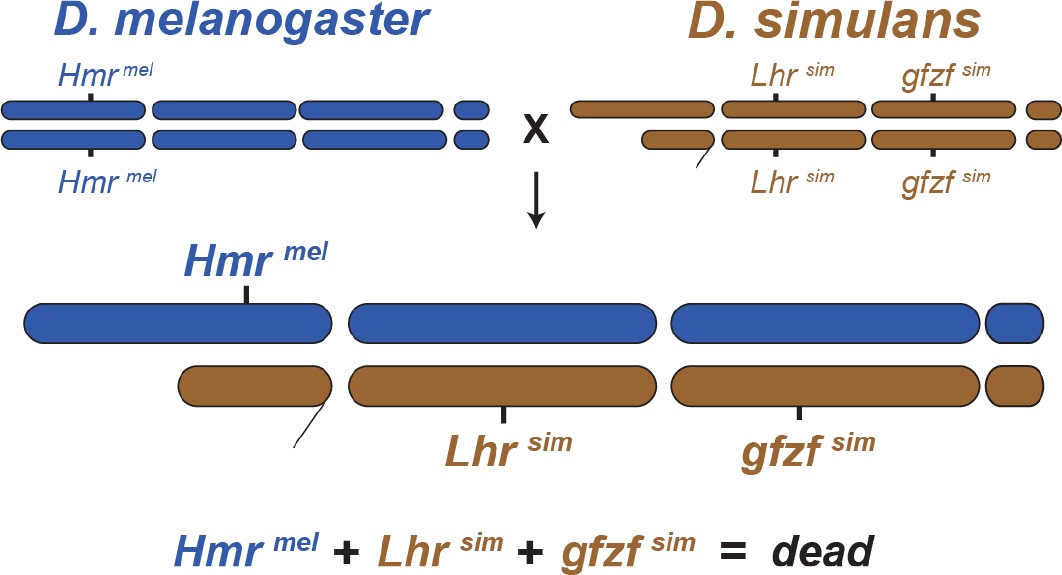
Three genes are required for F1 male lethality. *Hmr mel, Lhr sim,* and *gfzf sim* are all required for hybrid F1 male lethality between *D. melanogaster* and *D. simulans*.

*Hmr* and *Lhr* (also known as Heterochromatin Protein 3 (HP3)) are not essential for viability within species (Watanabe 1979; Hutter and Ashburner 1987). Biochemical studies have shown that HMR and LHR physically bind to each other and function together in a complex with Heterochromatin Protein 1 (HP1a) (Brideau *et al.* 2006; Thomae *et al.* 2013; Satyaki *et al.* 2014). Null mutants of *Hmr* and *Lhr* in *D. melanogaster* show defects in sister chromatid detachment during anaphase (Blum *et al.* 2017) and show de-repression of transcripts from several families of transposable elements and satellite DNA (Thomae *et al.* 2013; Satyaki *et al.* 2014). Both proteins are over-expressed in hybrid F1 males, and hybrids display a pattern of de-repression of transcripts from transposable elements and satellite DNA (Thomae *et al.* 2013; Satyaki *et al.* 2014). The molecular mechanism for how *Hmr* and *Lhr* interact with *gfzf* to form a lethal hybrid incompatibility remains unclear.

*gfzf* encodes a protein that contains 4 FLYWCH zinc finger domains, and a GST domain (Dai *et al.* 2004). In contrast to HMR and LHR, there is no evidence for a physical interaction of either protein with GFZF. Unlike *Hmr* and *Lhr*, *gfzf* is essential for viability in *D. melanogaster* (Provost *et al.* 2006). Loss of function mutants of *gfzf* die shortly after hatching into larvae, in a pattern reminiscent of hybrid F1 male lethality (Provost *et al.* 2006). *gfzf* has been identified repeatedly in several genetic screens, including those designed to identify suppressors of the *Killer-of-prune* system (Provost *et al.* 2006), cell-cycle regulation (Ambrus *et al.* 2009), DNA damage induced cell-cycle checkpoints (Kondo and Perrimon 2011), Ras/MAPK signaling (Ashton-Beaucage *et al.* 2014), and Polycomb complex regulation (Gonzalez *et al.* 2014). One potential explanation for the role of *gfzf* in such a variety of processes comes from a recent study where *gfzf* was identified as a transcriptional co-activator through an association with Motif1 binding protein (M1BP), and shown to bind the transcriptional start sites of genes relevant to the genetic screens where *gfzf* was identified as a hit (Baumann *et al.* 2018).

Here, we investigate the properties and functional differences of *gfzf* between *D. melanogaster* and *D. simulans*, and in F1 hybrids between these species. First, our cytological analyses show that there is little difference in protein localization of GFZF between the two species except at telomeric sequences. This differential binding of GFZF at telomeres persists in inter-species hybrids. These differences in localization appear to be due to changes in sequence composition at telomeres between *D. melanogaster* and *D. simulans,* and not due to species-specific binding by GFZF. Second, we investigate the pattern of GFZF and HMR co-localization, and find that the two proteins do not normally co-localize in *D. melanogaster*. In inter-species hybrids, however, HMR shows extensive mis-localization to GFZF sites, and this mis-localization requires the presence of *gfzf ^sim^*. Third, we find by ChIP-Seq that over-expression of HMR and LHR causes HMR to mis-localize to GFZF binding sites, indicating that HMR has some natural low affinity for GFZF sites. Finally, we conduct a proteomics analysis of GFZF and find several intriguing candidate *gfzf* interacting proteins that shed light on its role as a chromatin regulator, including the INO80 chromatin remodeling complex. Together, these results provide the first hints of the allele-specific properties of *gfzf*, and the novel molecular interactions between *Hmr* and *gfzf* in hybrids between *D. melanogaster* and *D. simulans*.

## Results

### Different localization patterns of GFZF at *D. melanogaster* and *D. simulans* telomeres

Though *gfzf* is a chromatin associated factor that has many binding sites in the genome (Baumann *et al.* 2018), it is not clear if this property is consistent between *D. melanogaster* and *D. simulans*. To determine whether *gfzf ^mel^* and *gfzf ^sim^* show differences in chromatin localization patterns, we developed an antibody that can recognize the GFZF protein from both species (Supplemental Figure 1). We performed immunostaining for natively expressed GFZF across multiple tissues, and found that it consistently localizes to the nucleus in *D. melanogaster* and *D. simulans* (Supplemental Figures S2 and 3). As we did not observe differences in the nuclear localization of GFZF at the cellular level, we next investigated the patterns of GFZF localization on polytene chromosomes. Consistent with previous reports, we found that GFZF localizes to many discrete bands on polytene chromosomes in *D. melanogaster* (Figure 2 A, B). However, we, also noticed strong localization of GFZF to the ends of several chromosomes, most notably the ends of the X and 2R chromosomes in *D. melanogaster*. These patterns are clearly visible both in our images and also in a re-examination of previously reported polytene analyses (Baumann *et al.* 2018).

**Figure 2.**
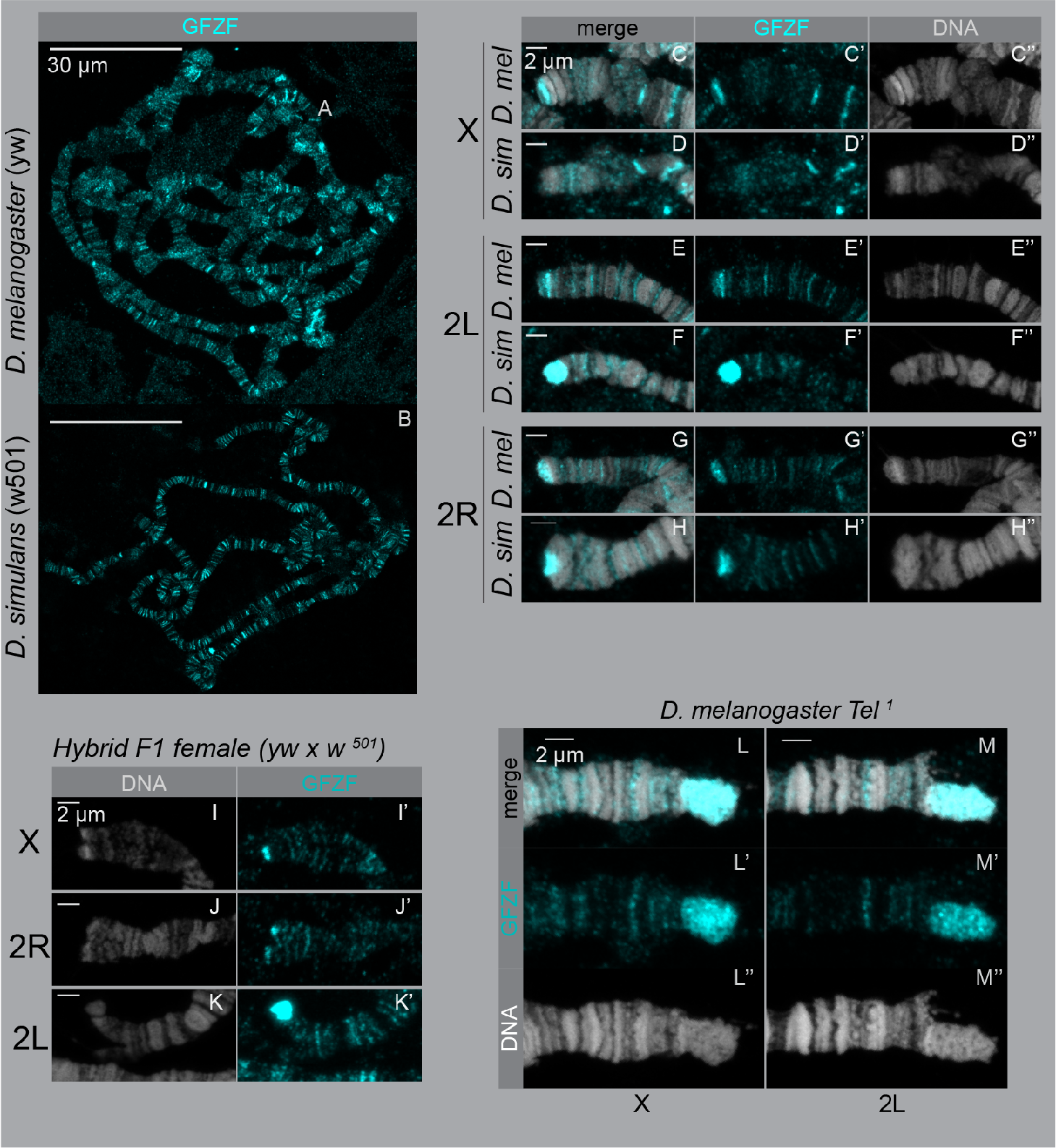
GFZF localization is different at chromosome ends in *D. melanogaster* and *D. simulans*. Polytene chromosomes were stained with anti-GFZF and Hoechst. A-B) Whole polytenes from *D. melanogaster* and *D. simulans*. In both species, GFZF localizes to multiple bands across the genome. C-H) Magnified ends of polytene chromosomes from *D. melanogaster* and *D. simulans*, stained for GFZF and DNA. I-K) Hybrid polytene chromosome ends of X, 2R, and 2L. There is a differential localization pattern on X and 2L, but not on 2R. H-I) GFZF binds telomere retrotransposon repeats. The *Tel ^1^* mutant has extended telomeres due to additional replication of the telomeric retrotransposons. In F1 progeny between *yw* and *Tel ^1^*, the GFZF signal extends with the elongated *Tel ^1^* telomere.

HMR localizes to the telomere capping complex (along with other locations in the genome) (Thomae *et al.* 2013; Satyaki *et al.* 2014). To test whether GFZF localizes to the telomere capping complex or the retrotransposon repeats that comprise the telomeres of *Drosophila*, we used a *Tel ^1^* mutant strain that has extended telomeric retrotransposon repeats (Siriaco *et al.* 2002). An increase of GFZF staining in *Tel ^1^* polytenes and the lack of co-localization with Heterochromatin Protein 1 indicate that GFZF localizes to the retrotransposon repeats at telomeres but not to the telomere cap (Figure 2 L, M, Supplemental Figure 4).

We further investigated the patterns of GFZF localization on polytene chromosomes in *D. simulans* and found that most bands of GFZF binding appear qualitatively similar between the two species. However, we uncovered striking species-specific differences in GFZF localization at telomeres (Figure 2 C-H). On the X chromosome, there is a bright band of staining on the end in *D. melanogaster* but not in *D. simulans*. This pattern is reversed at the end of 2L, where there is a bright patch of staining in *D. simulans* but not *D. melanogaster*. The end of chromosome arm 2R shows similar staining in both species, whereas the ends of 3L and 3R have little staining in either species. These patterns of species-specific GFZF staining at the ends of chromosomes seen in pure *D. melanogaster* and *D. simulans* individuals also persists in F1 hybrids between the two species (Figure 2 I-K, Supplemental Figure 5).

In F1 hybrids, GFZF localization on the *D. melanogaster* polytene chromosomes versus the *D. simulans* polytene chromosomes is dramatically different at telomeres, but otherwise appears identical at other regions of the genome. There is precedence for hybrid incompatibility proteins differentially binding DNA in hybrids, such as in the case of *Odysseus* which causes hybrid sterility in introgression males between *D. simulans* and *D. mauritiana* (Bayes and Malik 2009). Given the protein coding differences in GFZF between the two species, a simple hypothesis is that the two alleles of GFZF may show preferential binding to telomeres from its native species. To determine whether these patterns of differential localization reflect differences in binding specificities of the two alleles of GFZF or sequences composition differences at telomeres between the two species, we performed immuno-fluorescence studies using transgenes that carry GFP-tagged GFZF from either species. We found that both alleles of GFZF localize to telomeres in hybrids, with little or no allele-specific differences in their localization patterns (Figure 3). These results suggest that the differential binding patterns of GFZF at telomeres are not caused by differences in the binding-specificities of the two forms of GFZF, and instead reflect the rapid divergence of telomeric sequence composition between species (Danilevskaya *et al.* 1998; Anderson *et al.* 2008).

**Figure 3.**
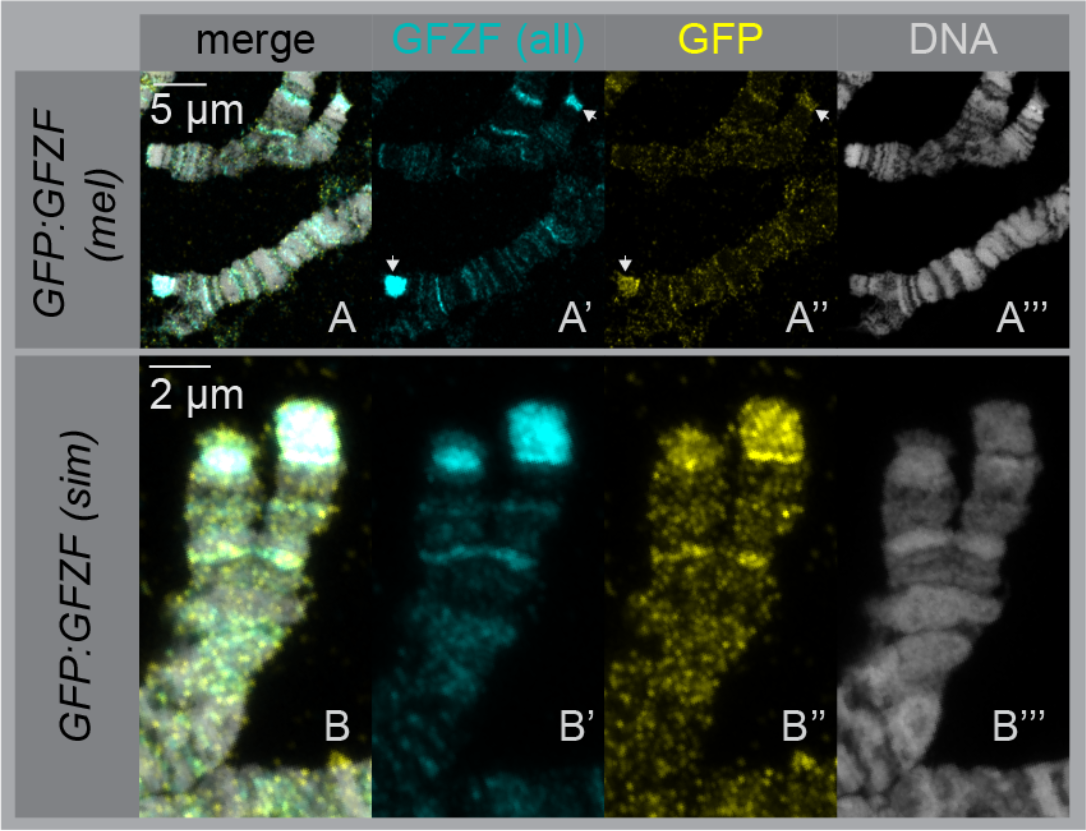
GFZF does not have species-specific binding patterns at polytene ends. F1 hybrid polytene chromosomes are stained for total GFZF and a EGFP-GFZF transgene from each individual species. In both cases, the anti-GFZF and anti-GFP signals co-localize, showing that there is not a difference in GFZF binding due to protein sequence.

### Altered localization of HMR in F1 hybrids requires GFZF^sim^

Our immunofluorescence studies with polytene chromosomes indicate that GFZF and HMR are in close proximity to each other at telomeres, where they localize to the telomeric retrotransposon repeats and the telomere capping complex respectively. To directly examine interactions between GFZF and HMR, we co-stained polytene chromosomes with antibodies for each protein. In *D. melanogaster*, we found that GFZF and HMR form separate bands through most of the genome, and rarely localize to the same band (Figure 4A, B). At telomeres, it appears that HMR localizes immediately distal to GFZF, consistent with the pattern that we observed with GFZF and HP1a. We attempted to detect the native localization of LHR in these contexts. While an LHR antibody exists that has been used previously for biochemical assays (Thomae *et al.* 2013), we failed to detect an LHR immunofluorescent signal in our experiments.

Previous observations show that HMR spreads to many additional sites in the polytene chromosomes of F1 hybrids (Thomae *et al.* 2013). We co-stained hybrid polytenes for GFZF and HMR and found that many of these new HMR chromatin sites co-localize with GFZF sites (Figure 4D). In euchromatic regions of F1 hybrids, there are more instances of HMR banding, where HMR appears to spread to many more locations throughout the genome. At the telomeres, HMR is no longer confined to the telomere cap, and instead leaks into the GFZF regions. Not all GFZF sites co-stain for HMR, suggesting that this overlap of HMR and GFZF localization in F1 hybrids is incomplete.

To test if the altered localization of HMR to GFZF sites in F1 hybrids is dependent on GFZF, we depleted allele specific versions of GFZF and measured HMR and GFZF co-localization. We used an Actin5C-GAL4 driver to knockdown *gfzf ^sim^* expression via RNAi, which has previously been shown to be sufficient to rescue hybrid male viability (Phadnis *et al.* 2015). Interestingly, reducing the expression of *gfzf ^sim^* greatly reduced the spreading of HMR into euchromatic regions of the genome, causing the HMR banding pattern to largely revert to the pattern seen in pure species individuals (Figure 4H, Supplemental Figure 6). We next tested whether a null allele of *Lhr ^sim^*, which is also sufficient to rescue hybrid male viability, similarly restored normal HMR localization. Here, we examined polytene chromosomes of hybrids from a cross between *melanogaster yw* females to *D. simulans Lhr^1^* males. In contrast to our results with reducing the expression of *gfzf ^sim^*, we find that the loss of function mutation in *Lhr ^sim^* does not prevent the spreading of HMR to the euchromatic regions of the genome (Figure 4E). Next, to test whether removing *gfzf ^sim^* versus removing *gfzf ^mel^* showed differences in their effect on HMR mis-localization, we crossed females carrying a null allele of *D. melanogaster gfzf* to *D. simulans* males, and found that removing GFZF ^mel^ does not stop the spreading of HMR (Figure 4F). We quantified these results by counting instances of overlapping peaks along polytene chromosomes, and found that reduction of GFZF ^sim^ is the only perturbation that significantly decreases the altered localization of HMR and to GFZF sites (Figure 4C). These results indicate that the mis-localization of HMR is dependent on GFZF ^sim^, but not on GFZF ^mel^ or LHR ^sim^.

**Figure 4.**
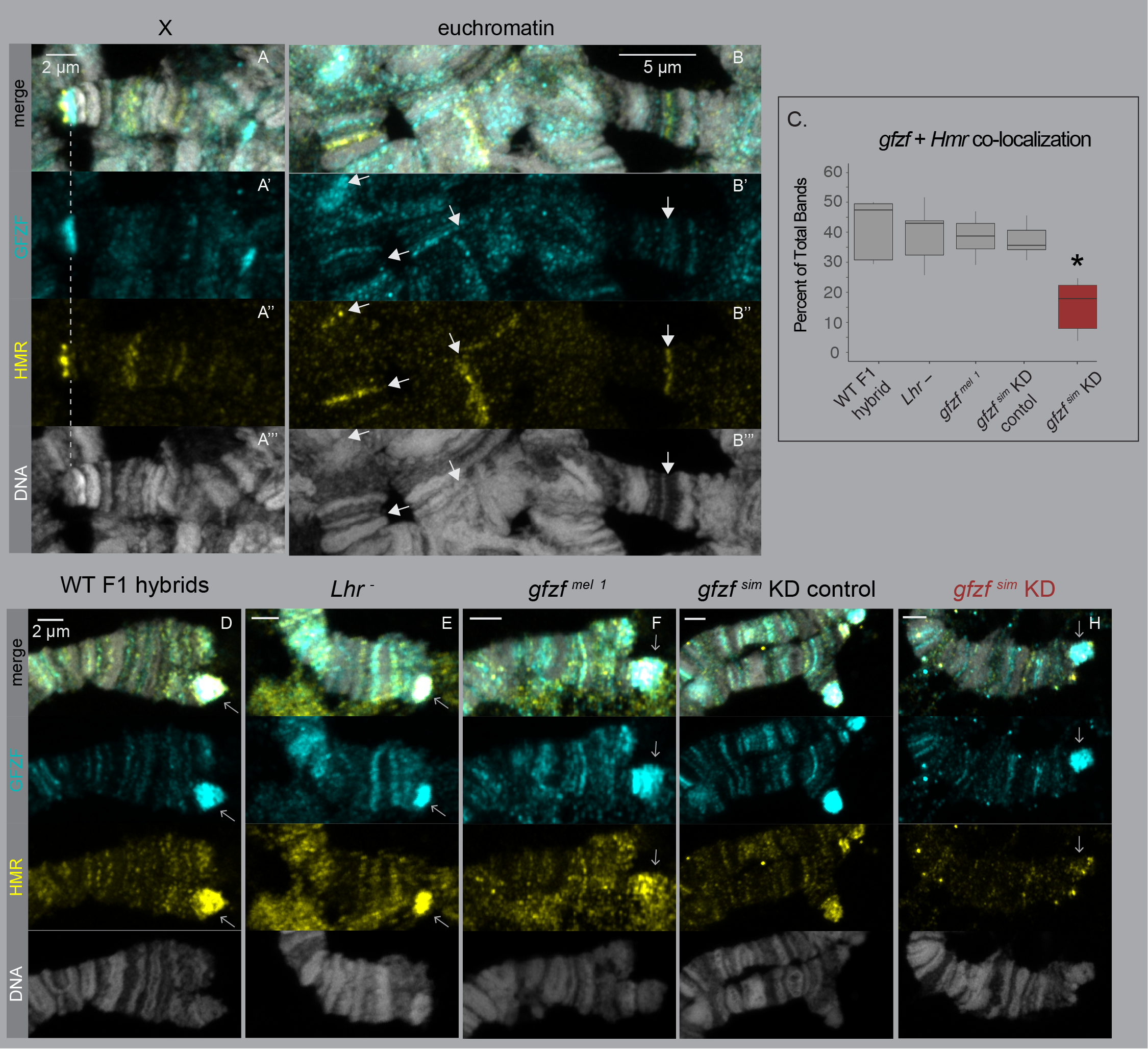
Aberrant co-localization of GFZF and HMR in F1 hybrids requires GFZF^sim^. A-B) F1 hybrid polytene chromosomes are stained with anti-GFZF and anti-HMR. There are few instances of co-localization, even when they are in close proximity at the telomeres. C) Quantification of co-localization. The number of HMR / GFZF co-occurring bands is presented as a percentage of the total bands recorded. * *p* < 0.0001 by a pairwise Wilcoxon rank sum test. D-H) Hybrid F1 female polytene chromosomes, stained for anti-GFZF and anti-HMR. All images show the telomeres of 2L. D) WT hybrids of *yw* and *w ^501^*. HMR spreads out to GFZF bands. E) Hybrids of *yw* crossed to *Lhr ^1^*. E) Hybrids of *gfzf ^2^* crossed to *w ^501^*. F) Hybrid of *gfzf ^1^* crossed to *w ^501^*. G-H) Hybrids from UAS-RNAi(*gfzf ^sim^*); Actin5C-GAL4 / CyO crossed to *w ^501^*. GAL4 vs CyO samples were identified by inversions on chromosome 2. In GAL4 expressing samples, HMR spreading is reduced and co-localization has decreased.

### Over-expressed HMR in *D. melanogaster* mis-localizes to GFZF sites without direct protein-protein interactions

We next asked if experimental manipulation within species could recapitulate an ectopic interaction between GFZF and HMR. A simple explanation for the mis-localization of HMR to GFZF binding sites in hybrids is that HMR may have some natural affinity for GFZF sites, but does not normally localize to these sites when *Hmr* is expressed at a low level. HMR is known to be over-expressed (along with LHR) in hybrids and, in this context, this over-expression of HMR may be responsible for its localization to GFZF binding sites. Alternatively, over-expressed HMR may form new protein-protein interactions with GFZF and localize to GFZF-binding sites. To test these ideas, we over-expressed HMR and LHR in *D. melanogaster* S2 cell culture, and performed ChIP-Seq with HMR to identify its genomic binding sites and compared them to endogenous binding sites. When both proteins are over-expressed, HMR localizes to new sites throughout the genome in a pattern consistent with our observations of HMR localization to GFZF binding sites in hybrids. These new sites overlap with GFZF binding sites, suggesting that HMR overexpression is sufficient to induce its mislocalization to GFZF binding sites (Figure 5, Supplemental Figure 8, 9).

**Figure 5.**
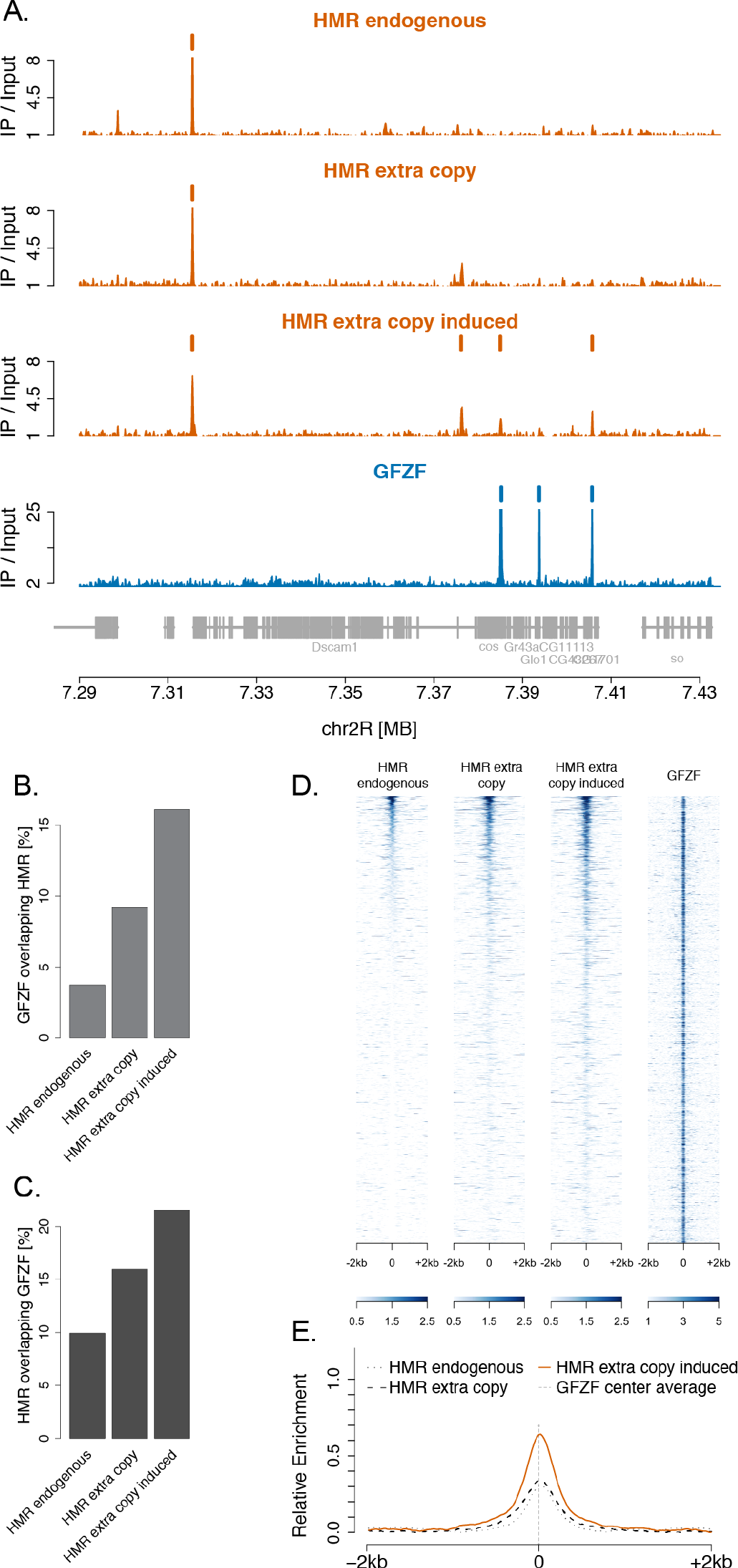
Over-expression of *Hmr* and *Lhr* in *D. melanogaster* causes HMR binding to GFZF genomic sites. Example of HMR and GFZF Chip-Seq plot over a region of chromosome 2R. HMR peaks are found coincident with GFZF peaks when HMR is over-expressed. B-C) The percentage of GFZF peaks that overlap HMR peaks and percentage of HMR peaks that overlap GFZF peaks increase genome wide with HMR over-expression. D) Heatmap for HMR enrichment at GFZF peaks in the three conditions, sorted by HMR signal strength in the induced condition. Across the three conditions, each line represents the same 4kb stretch around a GFZF peak. The GFZF heatmap shows the GFZF signal strength at each of these positions. E) Histogram of relative enrichment of HMR around GFZF peaks at all GFZF peaks for the three conditions.

In contrast to HMR and LHR, the protein interaction partners of GFZF remain unknown. As a result, we understand little about the molecular processes that GFZF is involved in. To identify protein complexes that stably interact with GFZF, we next performed AP-MS (affinity purification combined with Mass Spectrometry) experiments in embryonic nuclear extracts from the *D. melanogaster* Oregon-R strain under native conditions using our anti-GFZF antibody. Mass spectrometry analyses of the purified material uncovered interactions of GFZF in protein complexes involved in chromatin remodeling (INO80 complex), components of transcription complexes, and protein translation (Figure 6, Supplemental Figure 10). The interactions with the INO80 complex is particularly interesting because it plays an important role in chromatin remodeling, and is involved in various processes including DNA replication and repair (Clapier and Cairns 2009). The stable interactions of GFZF with these complexes provide important hints regarding the molecular processes that GFZF is involved in. Most notably, HMR and LHR are not present among the proteins that interact with GFZF. Consistent with previous results from AP-MS experiments with HMR (Thomae *et al.* 2013), these experiments further suggest that the mis-localization of HMR to GFZF binding sites is unlikely to be mediated by a direct interaction between these proteins.

**Figure 6.**
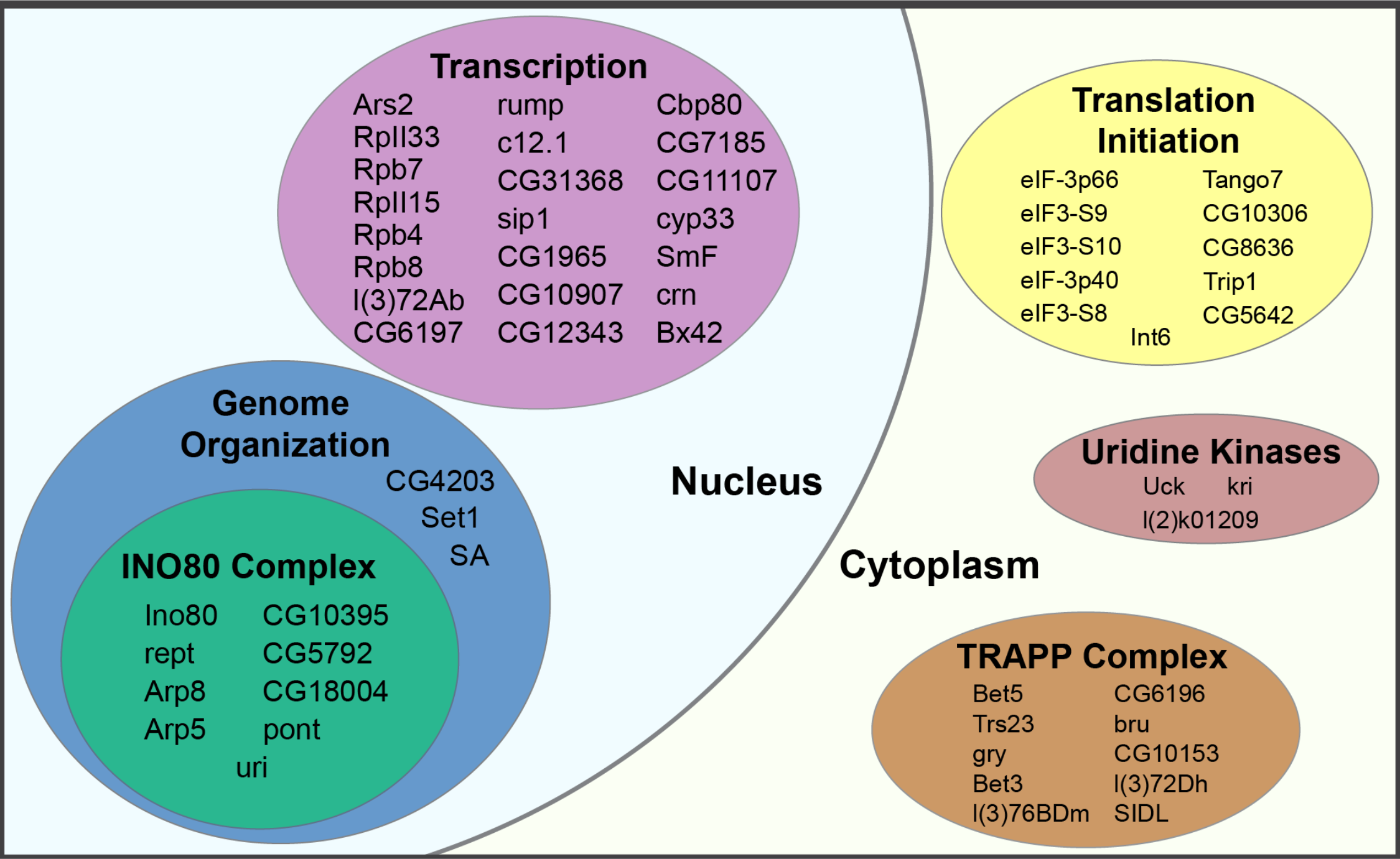
The proteome of GFZF. The proteome of GFZF by AP-MS. Major protein groups are shown in this figure – the full interactome is displayed in Supplemental Figure 10.

## Discussion

*Hmr*, *Lhr*, and *gfzf* genetically interact in a single hybrid male lethal incompatibility between *D. melanogaster* and *D. simulans* (Barbash *et al.* 2003; Brideau *et al.* 2006; Phadnis *et al.* 2015). All three genes evolve rapidly under recurrent positive selection and encode chromatin binding proteins. While HMR and LHR physically bind each other and function together in a single complex, the connection between either of these proteins to *gfzf* remains mysterious. Our results indicate that GFZF localizes to discrete genomic locations, but the binding profiles of GFZF do not overlap with those of HMR. These proteins come close to each other at chromosome ends, where GFZF is heavily enriched at the retrotransposon arrays that form fly telomeres and HMR/LHR function together in the telomere capping complex. Despite this close proximity of GFZF with HMR at chromosome ends, GFZF does not overlap with HMR and instead the two proteins reflect the boundary between the retrotransposon arrays at telomeres and the telomere capping complex.

However, in inter-species hybrids these proteins meet each other in the form of altered chromatin localization of HMR to GFZF binding sites. This change is easily observed in the form of HMR localization to the retrotransposon arrays where GFZF is present, and also at other euchromatic GFZF-binding sites. Previously, HMR has been observed to localize to additional sites in hybrids compared to its within species binding patterns, but the nature of these sites remained unknown (Thomae *et al.* 2013). These additional sites can now at least partially be accounted for through GFZF-binding sites. Our results show that reducing the expression of the incompatible *gfzf ^sim^* allele restores the normal localization patterns of HMR. Although removing the *D. simulans* alleles of either *Lhr* or *gfzf* rescues hybrid male viability, removing *Lhr ^sim^* does not restore the normal localization pattern of HMR indicating distinct roles for *Lhr* and *gfzf* in the hybrid incompatibility. Together, these results provide evidence for a new molecular interaction between the hybrid incompatibility factors involved in hybrid male lethality between *D. melanogaster* and *D. simulans*.

The observation that reducing *gfzf ^sim^* restores normal localization of HMR raises the possibility that a novel physical interaction between GFZF ^sim^ and HMR may underlie the altered localization of HMR to GFZF binding sites in hybrids. Our proteomic analyses suggest that at least within *D. melanogaster*, no such physical interactions normally exist between these proteins. Similar proteomic analyses in inter-species hybrids are technically challenging. However, our experiments with *D. melanogaster* cultured cells show that over-expressing *Hmr* and *Lhr* is sufficient to localize HMR to GFZF binding sites in the absence of *gfzf ^sim^*. A novel physical interaction between HMR and GFZF may, therefore, not be necessary for the altered localization patterns seen in hybrids. Instead, a natural affinity of surplus HMR for GFZF binding sites may be sufficient to explain the altered binding patterns observed in hybrids. *Hmr* and *Lhr* are known to be over-expressed in hybrids between *D. melanogaster* and *D. simulans* (Thomae *et al.* 2013). Consistent with these observations, our results suggest a model where *gfzf ^sim^* may be responsible for the over-expression of *Hmr* and *Lhr*, which in turn leads to the leakage of surplus HMR to GFZF binding sites in hybrids.

The only obvious difference in GFZF localization between *D. melanogaster* and *D. simulans* using polytene analyses is at their telomeres. The *D. melanogaster* and *D. simulans* alleles of GFZF, however, are capable of binding to the telomeric retro-transposon arrays at telomeres from either species. The differences in GFZF binding patterns between *D. melanogaster* and *D. simulans* thus involve little or no species-specific binding by the alleles of GFZF. Instead, these results reveal the evolution of dramatic copy number differences in the telomeric retro-transposon arrays between the two species. Interestingly, our proteomic analysis reveals a strong interaction between GFZF and nearly the entire INO80 complex. The INO80 complex is known to regulate telomere length and structure in yeast and in mice (Yu *et al.* 2007; Min *et al.* 2013). In addition, the *Tel^1^* mutation in *D. melanogaster* that results in the rapid expansion of telomeric retro-transposon arrays has also been mapped to the *ino80* locus (Reddy *et al.* 2018). The INO80 complex, thus, appears to be involved in the regulation of telomeres in both yeast and flies despite their dramatically different mechanisms of telomere maintenance. Further experiments to explore the role of GFZF in telomere regulation may help to determine how GFZF operates as a transcriptional co-activator, but also localizes to the highly repressed retrotransposon repeats of the telomere.

Although hybrids between *D. melanogaster* and *D. simulans* show increased expression of transposable elements including those at telomeres, this is not thought to be directly responsible for hybrid male lethality (Thomae *et al.* 2013; Satyaki *et al.* 2014). However, the need to control telomeric retrotransposons may have provided the substrate for the rapid evolution of *Hmr*, *Lhr* and *gfzf*. Telomeric retrotransposons must replicate for cells to live, yet rampant replication may cause widescale genome instability. Therefore, the regulation of copy number and the telomere/euchromatin boundary might be important stages of intragenomic conflict. All three hybrid incompatibly genes are positioned to be involved in both conflicts – in hybrids, the extension of HMR into the GFZF regions of telomeres could may reflect the loss of a boundary between the two regions. These ideas provide guidance for further experiments into understanding the functional consequences of the altered localization of HMR to GFZF binding sites, particularly at telomeres. Together, our results suggest that a deeper understanding of the molecular interactions of *gfzf* may shed light on telomere length regulation through retro-transposon copy number control, and its relation to the evolution and molecular mechanisms of hybrid incompatibilities.

## Materials and Methods

### Drosophila Husbandry and Strains

To produce hybrids between *D. melanogaster* and *D. simulans*, we set crosses of 3-4 virgin female *D. melanogaster* and 7-9 two day old *D. simulans* males. We allowed mating to occur at 25°C for 2-3 days, and then transferred the vials with progeny to 18°C to develop. Unless stated otherwise, our standard *D. melanogaster* line was *yw*, and our standard *D. simulans* line was *w ^501^*. For our polytene analyses, we set up all crosses with 3-4 males and females at 25°C for two days and reared the progeny 18°C until large 3^rd^ instar larvae were ready for dissection.

### Transgenic Flies

To construct flies that expressed EGFP.GFZF, we cloned the *gfzf* locus from both *D. melanogaster* and *D. simulans* into plasmid vectors for transformation into fly lines and created stable stocks. Briefly, we amplified 1KB upstream of the transcript start site of *gfzf* and 1KB downstream of the stop codon of *gfzf* in order to capture any local regulatory elements. We used Gibson Assembly (Gibson *et al.* 2009) to transform this segment of the *gfzf* locus with the sequence for EGFP (fused to the N-terminus of *gfzf* with a short linker sequence) into the pattB and pCasper4 vectors. Injections of these vectors into PBac{y[+]-attP-3B}VK00002 (Bloomington Stock 9723) by phiC31 transgenesis for *D. melanogaster* and *w^501^* by P-element mediated transgenesis for *D. simulans* were performed by The Best Gene Inc. We confirmed the successful integration of these vectors via a *mW* marker, and the expression of EGFP.GFZF by wide-field fluorescence microscopy. In all cases we found that the 1KB upstream and downstream regulatory regions were sufficient alone to drive expression of EGFP.GFZF at a moderate level in every cell type we examined (as compared with the native fluorescence of other transgenes we have used in our lab). All sequence information is available in the Supplemental Text (Supplemental Table 2). The UAS.*gfzf ^sim^* RNAi strain was developed for a previous study (Phadnis *et al.* 2015), and uses a shRNA construct to target a 6bp deletion that is a fixed difference between *D. melanogaster* and *D. simulans gfzf*.

### Polytene chromosome analysis

We isolated polytene chromosomes from the salivary glands of 3^rd^ instar larvae. We extracted the salivary glands into PBS and fixed them in 45% acetic acid 4% paraformaldehyde for 30 seconds. We then softened the salivary tissue in 22.5% acetic acid and 50% lactic acid for 1 minute. We flattened the glands and cleared tissue under a wide field dissecting scope between a coverslip and a glass slide. We fixed the tissues to the slide by freezing the sample with liquid nitrogen and removing the coverslip using a swift flicking motion. We stored samples in PBS at 4ºC for no more than 3 hours before proceeding to immunofluorescent antibody staining.

### Antibodies and Immunofluorescence

To fluorescently stain polytene chromosomes, we first washed them with PBS with 0.3% TritonX-100 (PBT). We then blocked them for 10 minutes with normal goat serum, and incubated them in with a primary antibody solution for 1hr and 20min at room temperature. The rabbit anti-GFZF primary antibody was developed by Covance against the target RRDVAEPAKGAQPDC. For our primary antibody concentrations, we used the following: rabbit anti-GFZF 1:200, rat anti-HMR 1:20 (Thomae *et al.* 2013), mouse anti-HP1 (DSHB C1A9), and chicken anti-GFP 1:200 (Abcam 13970). We washed the samples 6 times with PBT before incubating them in a secondary antibody solution for 1hr 20min at room temperature. All secondary antibodies were generated from goats and used at a concentration of 1:1000. We washed away the secondary antibody with three washes of PBT. To stain DNA, we incubated the samples with Hoescht 32258 (Thermo Fisher) at 1:1000 in PBT for 4 minutes. We washed twice more with PBS before mounting the samples under glass coverslips in Fluoromount-G. We imaged all our samples using the Ziess LSM880 Airyscan system, and processed all images using the Zen software from Ziess and the Fiji package in ImageJ.

### Co-localization analysis

To analyze the co-localization of HMR and GFZF in our polytene images, we measured the frequency of co-occurring bands. To do this, we drew a segmented line along DNA using tools in Fiji (a package manager for ImageJ), and generated a table of the intensities of the HMR and GFZF signal from their respective channels. To pick the location of our lines, we avoided regions that contained large, continuous stretches of signal from either channel, as to not artificially inflate the amount of co-localization. We also picked segments that contained at least one prominent band from both channels to ensure that signals could be clearly separated from background noise. We used the DNA channel to find continuous segments of DNA. For each condition, we measured at least five polytene preps from separate larvae. For each polytene, we made three measurements on different segments of the polytene and averaged the three measurements. To process the data, we wrote a script in python3 to detect local maxima (peaks in fluorescent intensity that represent bands) and count the co-occurrence of maxima in both samples vs the total number of maxima observed (Supplemental Figure 7). We avoided double-counting by setting a minimum distance between local maxima (0.6µM), and set a background threshold as 20% of the maximum intensity value to avoid noise in the fluorescent signal. We tested for statistical difference in our samples using by implementing pairwise comparisons of a Wilcoxon Rank Sum Test in R.

### Cell culture

*Drosophila* SL2 cells stably transfected with FLAG-HA-HMR and Myc-LHR under a CuSO4 inducible promoter (pMT) were grown at 26°C in Schneider *Drosophila* medium (Invitrogen) supplemented with 10% fetal calf serum and antibiotics (100 units/mL penicillin and 100 μg/mL streptomycin). Transfected cells were selected with 20 μg/mL Hygromycin B. Cells were grown to confluence in 550 mL flasks and induced for 20-24h with 250 μM CuSO4 before harvesting for chromatin immunoprecipitation.

### Embryo nuclear extraction for AP-Mass Spectrometry

*D. melanogaster* Oregon R flies were grown at 26°C with 50% humidity. Embryo collection and extraction were performed as in Kunert and Brehm 2008 (Kunert and Brehm 2008). Embryo nuclear extracts were used for immunoprecipitation coupled to mass spectrometry.

### Immunoprecipitation

Nuclear extracts from 0-12h Oregon R embryos were subjected to immunoprecipitation as follows. α-FLAG immunoprecipitation was performed using 20 μL of packed agarose-conjugated mouse α-FLAG antibody (ANTI-FLAG M2 Affinity gel, A2220 Sigma-Aldrich). α-GFZF IP was performed using 12 μL of rabbit antibody non-covalently coupled to 30 uL of Protein A/G Sepharose. Unspecific rat IgG non-covalently coupled to 30 μL of Protein A/G Sepharose through a bridging rabbit anti-rat IgG (Dianova, 312-005-046) and Protein A/G Sepharose alone (beads-only) were used as mock controls, in mass-spectrometry and immunoblots experiments, respectively. The steps that follow were the same for all the immunoprecipitations and were all performed at 4°C. The antibodies coupled with the solid phase were washed three times with IP buffer (25mM Hepes pH 7.6, 150 mM NaCl, 12.5 mM MgCl_2_, 10% Glycerol, 0.5 mM EGTA) prior to immunoprecipitation. Nuclear extracts were treated with benzonase (MERCK 1.01654.0001) and rotated 1h end-over-end at 4°C to digest nucleic acids. To remove insoluble material the digested extracts were centrifuged for 10’ at 20000 × g and the supernatant was transferred to new tubes containing the antibody-beads solution (IP buffer complemented with a cocktail of inhibitors containing Aprotinin, Pepstatin, Leupeptin, 0.25 μg/mL MG132, 0.2 mM PMSF, 1 mM DTT was added up to a total volume of 500 μL) and end-over-end rotated for 2h (α-FLAG) or 4h (α-GFZF and IgG). After incubation the beads were centrifuged at 400 × g and washed 3 times in IP buffer complemented with inhibitors and 3 additional times with NH_4_HCO_3_ before in beads digestion.

### Sample preparation for mass spectrometry

The pulled-down material was released from the beads by pre-digesting for 30 minutes on a shaker (1400 rpm) at 25°C with trypsin 10 ng/μL in 100 μL of digestion buffer (1M Urea, 50 mM NH_4_HCO_3_). After centrifugation the peptides-containing supernatant was transferred to new tubes and two additional washes of the beads were performed with 50 μL of 50 mM NH_4_HCO_3_ to improve recovery. 100 mM DTT was added to the solution to reduce disulphide bonds and the samples were further digested overnight at 25°C while shaking at 500 rpm. The free sulfhydryl groups were then alkylated by adding iodoacetamide 12 mg/mL and incubating 30 minutes in the dark at 25°C. Finally, the light-shield was removed and the samples were treated with 100 mM DTT and incubated for 10 minutes at 25°C. The digested peptide solution was then brought to a pH ≈ 2 by adding 4μL of trifluoroacetic acid (TFA) and stored at −20°C until desalting. Desalting was done by binding to C18 stage tips and eluting with elution solution (30% methanol, 40% acetonitrile, 0.1% formic acid). The peptide mixtures were dried and resuspended in 20 μL of formic acid 0.1% before injection.

### Sample analysis by mass spectrometry

Peptide mixtures were subjected to nanoRP-LC-MS/MS analysis on an Ultimate 3000 nano chromatography system coupled to a QExactive HF mass spectrometer (both Thermo Fisher Scientific) in 5 uL. The samples were directly injected in 0.1% formic acid onto the separating column (120 × 0.075 mm, in house packed with ReprosilAQ-C18, Dr. Maisch GmbH, 2.4 µm) at a flow rate of 0.3 µl/min. The peptides were separated by a linear gradient from 3% ACN to 40% ACN in 50 min. The outlet of the column served as electrospray ionization emitter to transfer the peptide ions directly into the mass spectrometer. The QExactive was operated in a TOP10 duty cycle, detecting intact peptide ion in positive ion mode in the initial survey scan at 60,000 resolution and selecting up to 10 precursors per cycle for individual fragmentation analysis. Therefore, precursor ions with charge state between 2+ and 5+ were isolated in a 2 Da window and subjected to higher-energy collisional fragmentation in the HCD-Trap. After MS/MS acquisition precursors were excluded from MS/MS analysis for 20 seconds to reduce data redundancy. Siloxane signals were used for internal calibration of mass spectra.

### Proteomics Data analysis

For protein identification, the raw data were analyzed with the Andromeda algorithm of the MaxQuant package (version 1.6.0.16) against the Flybase reference database dmel-all-translation-r6.12.fasta including reverse sequences and contaminants. Default settings were used except for: Variable modifications = Oxidation (M); Unique and razor, Min. peptides = 1; Match between windows = 0.8 min. iBAQ values were log2 transformed, imputed using the impute.MinProb function (q = 0.05, tune.sigma = 1) from the imputeLCMD R package (v2.0) and normalized by centering each sample to the median matched to the grand median. Statistical tests were performed by fitting a linear model and applying empirical Bayes moderation using the limma package (v3.34.9).

### ChIP-Seq

Chromatin immunoprecipitation was essentially performed as in Gerland et al., 2017 (Gerland *et al.* 2017). For each ChIP reaction, chromatin isolated from 1–2 × 10^6^ cells was incubated with rat anti-HMR 2C10 antibody pre-coupled to Protein A/G Sepharose through a rabbit IgG anti-rat. The samples were single-end, 50 bp sequenced with the Illumina HiSeq2000. An overview of all ChIP-Seq samples used, a list of HMR peaks used for further analyses is as well as all sequencing data are publicly available as described below. The raw reads were aligned to the *D. melanogaster* genome assembly (UCSC dm6) using Bowtie2 (2.2.9) and filtered for uniquely mapped reads using samtools (1.3.1) (Li et al., 2009). Input and sequencing depth normalized tracks were generated by Homer (4.9) and visualized using R graphics. Peak calling was performed using HOMER 4.9 with parameters – style factor -F 2 -size 200 for HMR and -style factor -F 6-L 6 -size 200 for GFZF. Downstream analysis steps were performed using R. The endogenous HMR binding data set (GSE86106, Gerland et al., 2017) and the GFZF binding data set (GSE105009, Baumann et al., 2017) are derived from NCBI GEO.

### Data access

All proteomics data are available in the Project PXD010712 in the PRIDE archive of EMBL-EBI. All ChIP-sequencing data are available in the Project GSE118291 in the GEO archive of NCBI.

## Acknowledgements

We thank A. Thomae for the cell lines, N. Prayitno for sharing some embryo extracts, T. Gerland and P. Krueger for help with ChIP-Seq and I. Forne and A. Schmidt for measuring MS samples. We thank M. Levine, H.S. Malik, and S. Gross for their helpful comments. This work was supported by the National Institutes of Health (Developmental Biology Training Grant 5T32 HD0741 (JCC); R01 GM115914 (NP), a Mario Capecchi endowed chair (NP), and the Pew Biomedical Scholars Program (NP), a predoctoral grant to A.L (QBM) and a grant from the Deutsche Forschungsgemeinschaft DFG (CIPSM) (AI). We also thank S. Phadnis for continued moral support.

---

**Supplemental Figure 1.**
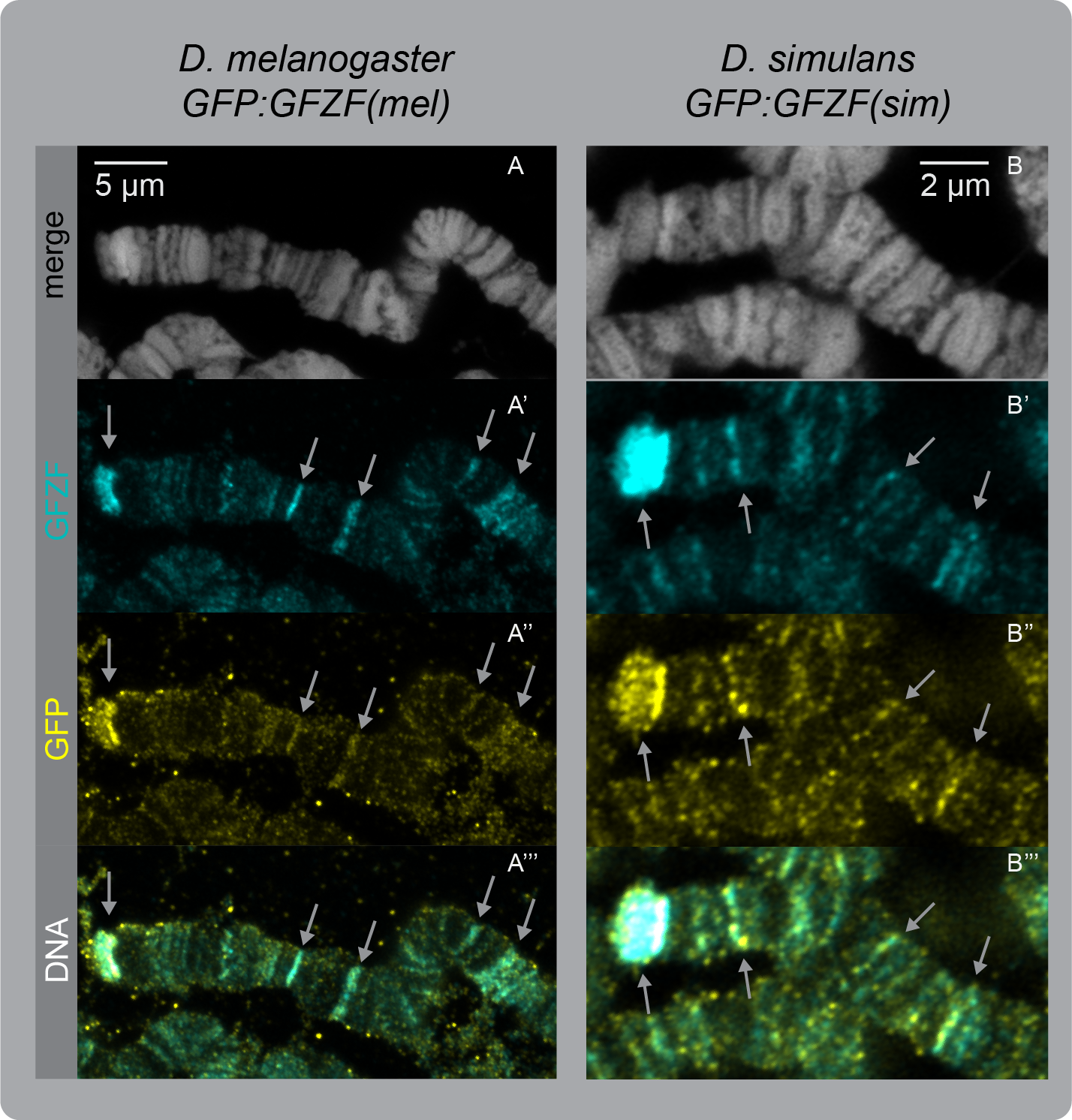
GFZF antibody binds GFZF ^mel^ and GFZF ^sim^. Polytene chromosomes from transgenic lines with EGFP.GFZF ^mel^ and EGFP.GFZF ^sim^ co-stained with anti-GFP and anti-GFZF antibodies. White arrows point to strong bands of co-localization.

**Supplemental Figure 2.**
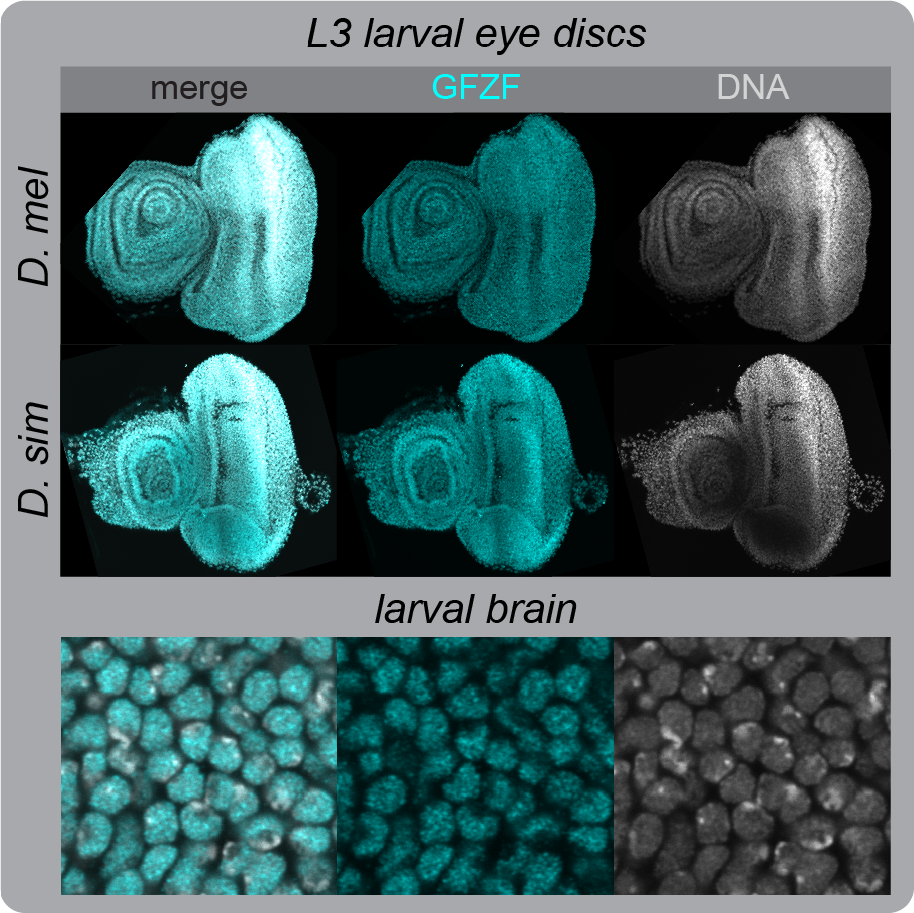
GFZF is expressed in the nucleus during development in *D. melanogaster* and *D. simulans*. Larval eye discs stained with anti-GFZF and Hoechst from *D. melanogaster* and *D. simulans*. Mid panels are zoomed in, single Z slices from the eye disc samples. In all these samples, GFZF co-localizes with Hoechst.

**Supplemental Figure 3.**
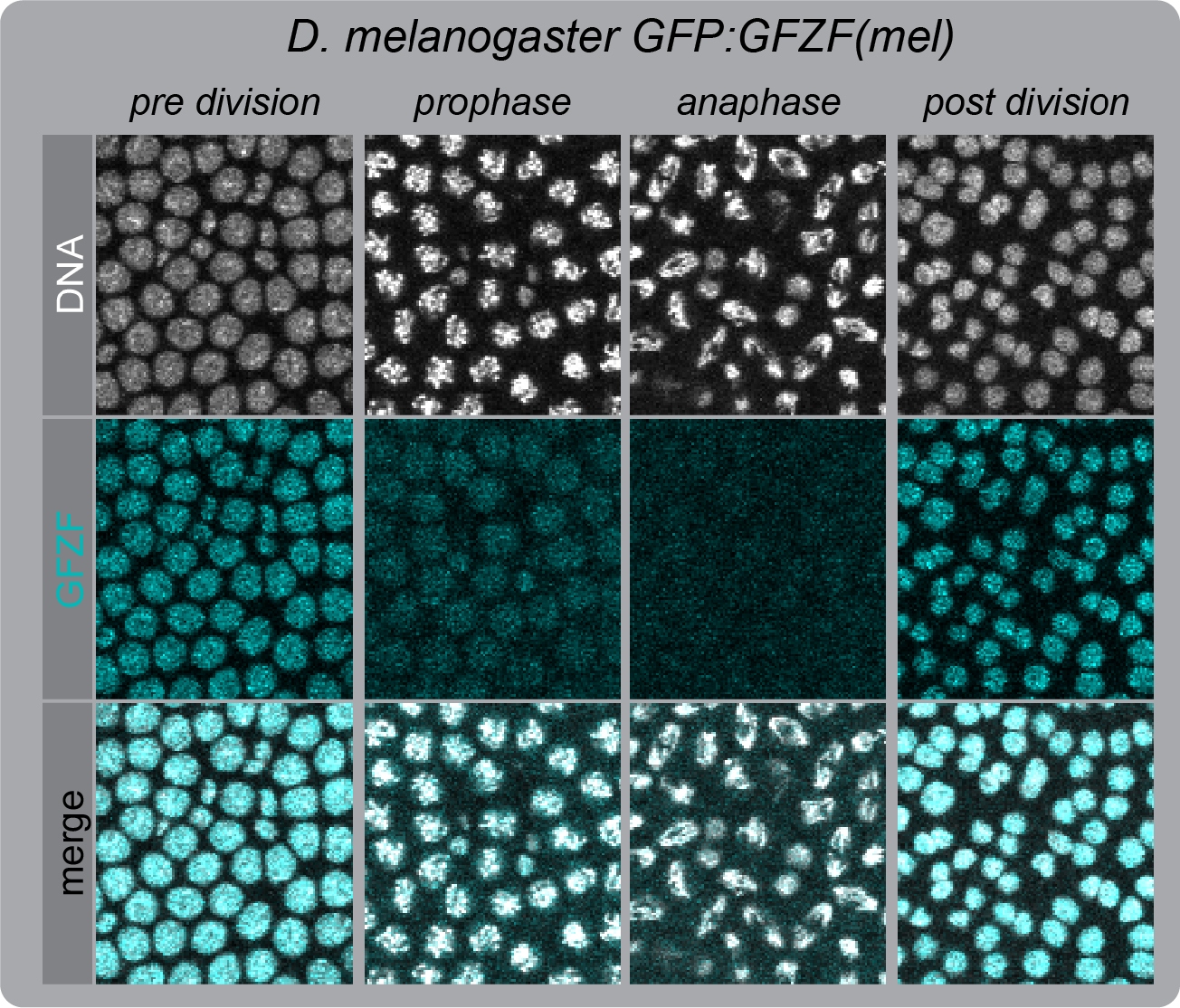
GFZF is absent from the chromatin of dividing cells. Live embryo imaging of EGPF.GFZF; H2av.RFP in *D. melanogaster* over the stage 6 – stage 7 cell division. Images were taken at 20 second intervals, and the panels here are representative of the cell cycle stages indicated in the legend. During mitosis, the EGFP signal depletes from the nucleus as chromatin condenses and is almost absent at the onset of anaphase. The signal returns early in the next prophase, and the process repeats.

**Supplemental Figure 4.**
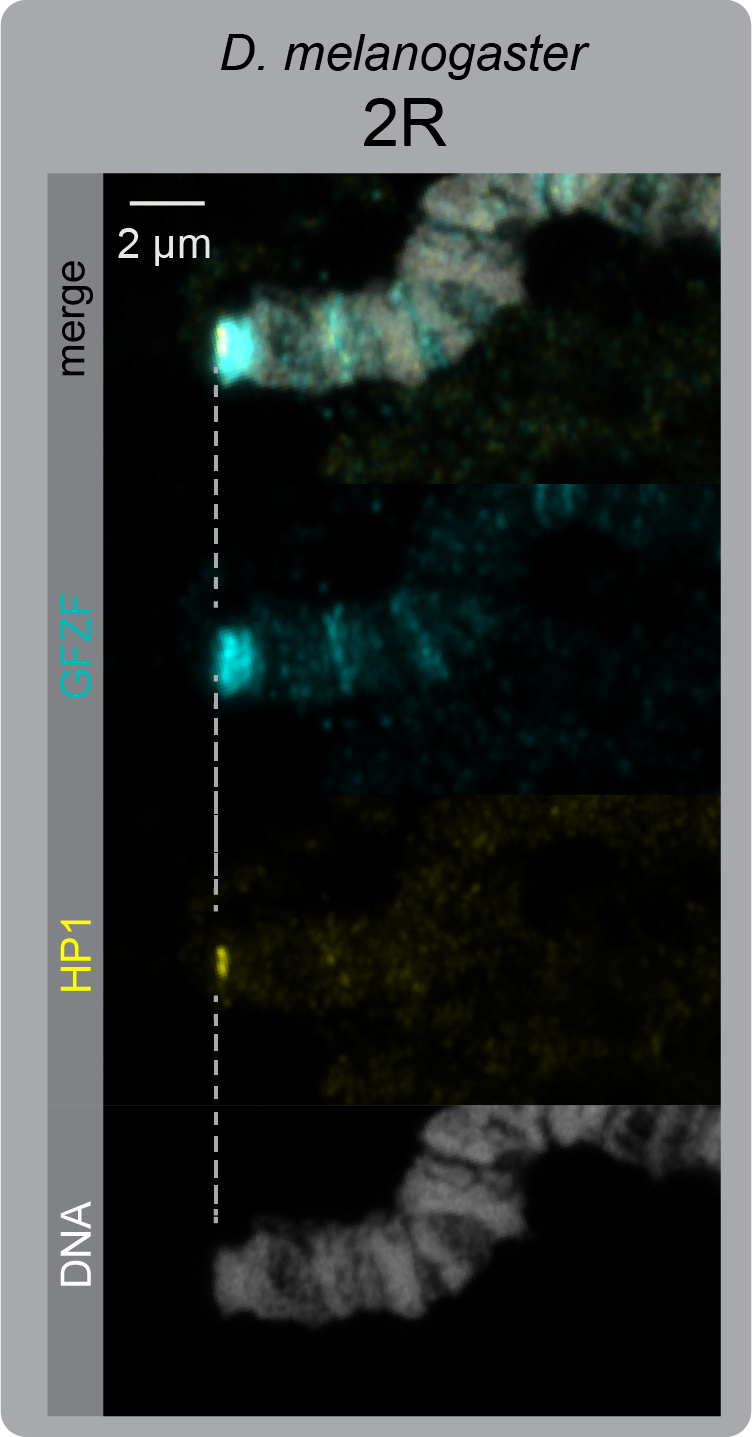
GFZF and HP1a do not co-localize at the ends of chromosomes, indicating that GFZF does not bind the telomere cap. Co-staining for anti-GFZF and anti-HP1a. The alignment via the dashed line shows that the HP1a signal is just distal of the GFZF signal, similar to the pattern that we observe with GFZF and HMR.

**Supplemental Figure 5.**
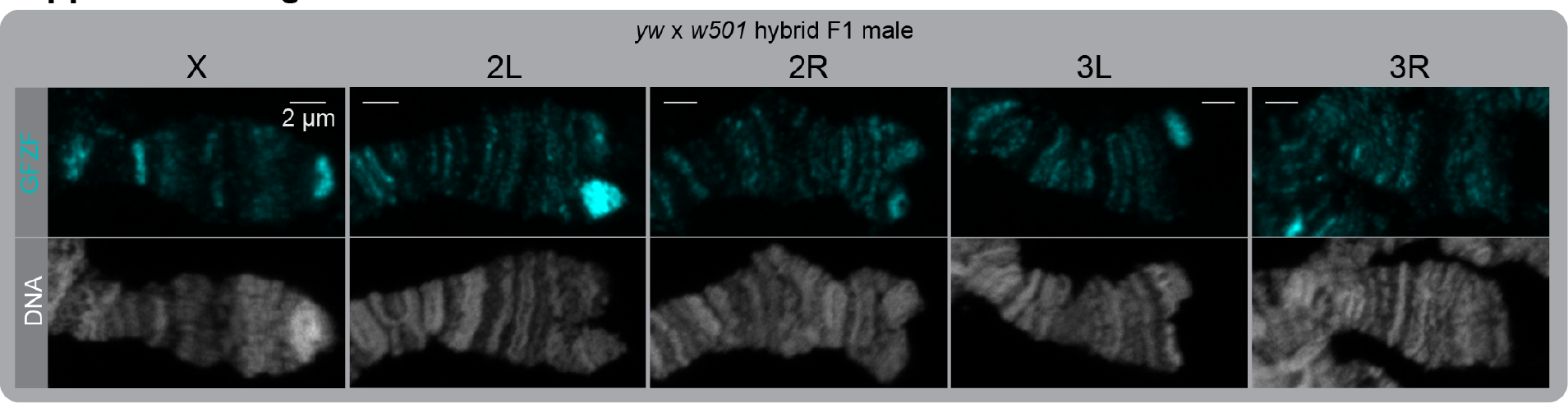
GFZF localization at telomeres is different in hybrid F1 males. We analyzed polytene chromosomes from rare hybrid males (~5%) that grow to be large 3^rd^ instar larvae, and found that GFZF has staining patterns similar to that of hybrid F1 females. Though these males grow, they do not become viable adults as indicated by the lack of adult males recovered. We confirmed that these are true hybrid males by two methods; the *yw D. melanogaster* X chromosome causes larval mouth hooks to be lighter in color, and the polytene karyotype contained only 1 X that stained for GFZF at the telomere, indicating that it is in fact the *D. melanogaster* X.

**Supplemental Figure 6.**
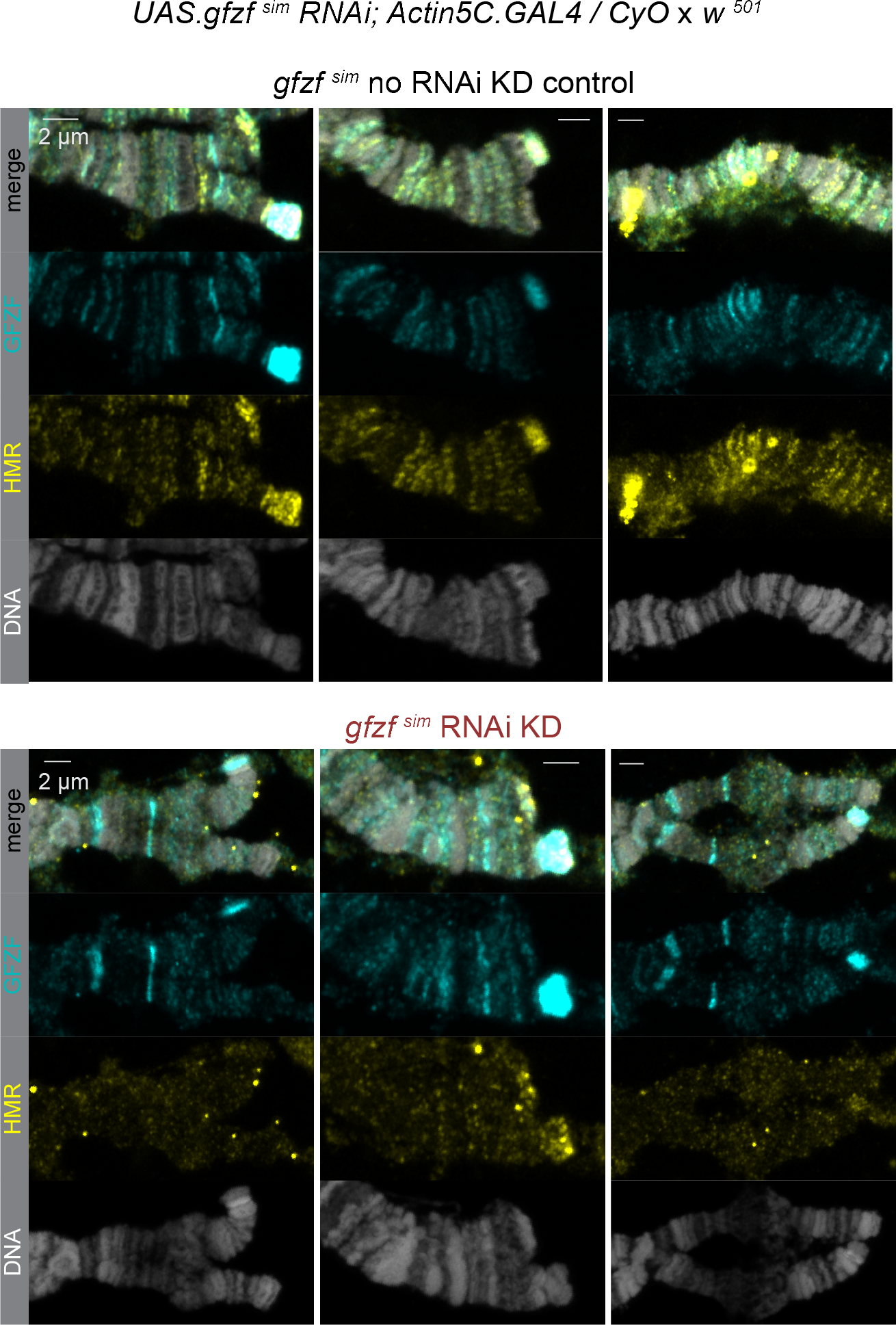
Additional images of *gfzf ^sim^* knockdown hybrids. The top panels hybrid polytenes from control GAL4-samples. The bottom panels are hybrid polytenes are from GAL4+ *gfzf ^sim^* knockdown samples. HMR staining and HMR / GFZF co-localization is reduced in the *gfzf ^sim^* knockdown samples.

**Supplemental Figure 7.**
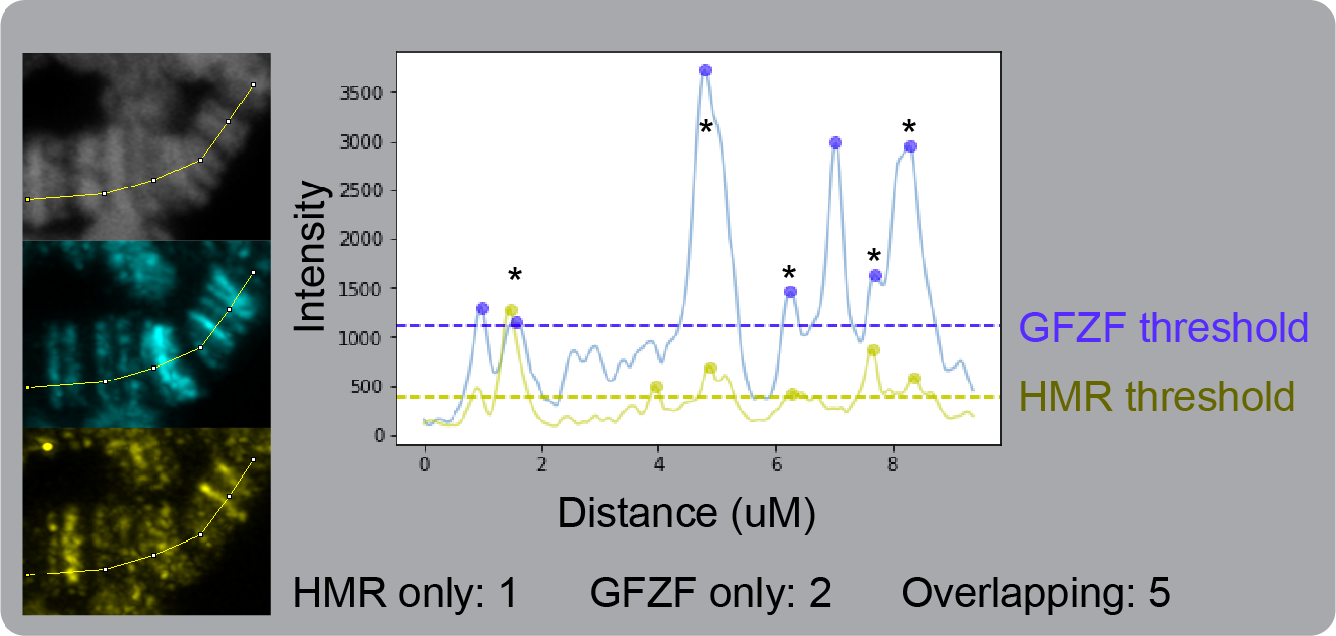
Example of GFZF / HMR overlap calculation process. Polytene fluorescent images analyzed for co-localization between GFZF and HMR along the DNA axis. The sample shown here is only several microns for sake if clarity – for our data presented in figure 4, we mapped ~150 microns of each polytene. A) We used the segmented line tool in Fiji to trace the DNA line of the polytene chromosomes, and created fluorescent intensity outputs for the GFZF and HMR channels separately. B) We determined local maxima for GFZF and HMR. C) Tabulated the peaks of each channel individually and the peaks that were present in both channels. More details in methods.

**Supplemental Figure 8.**
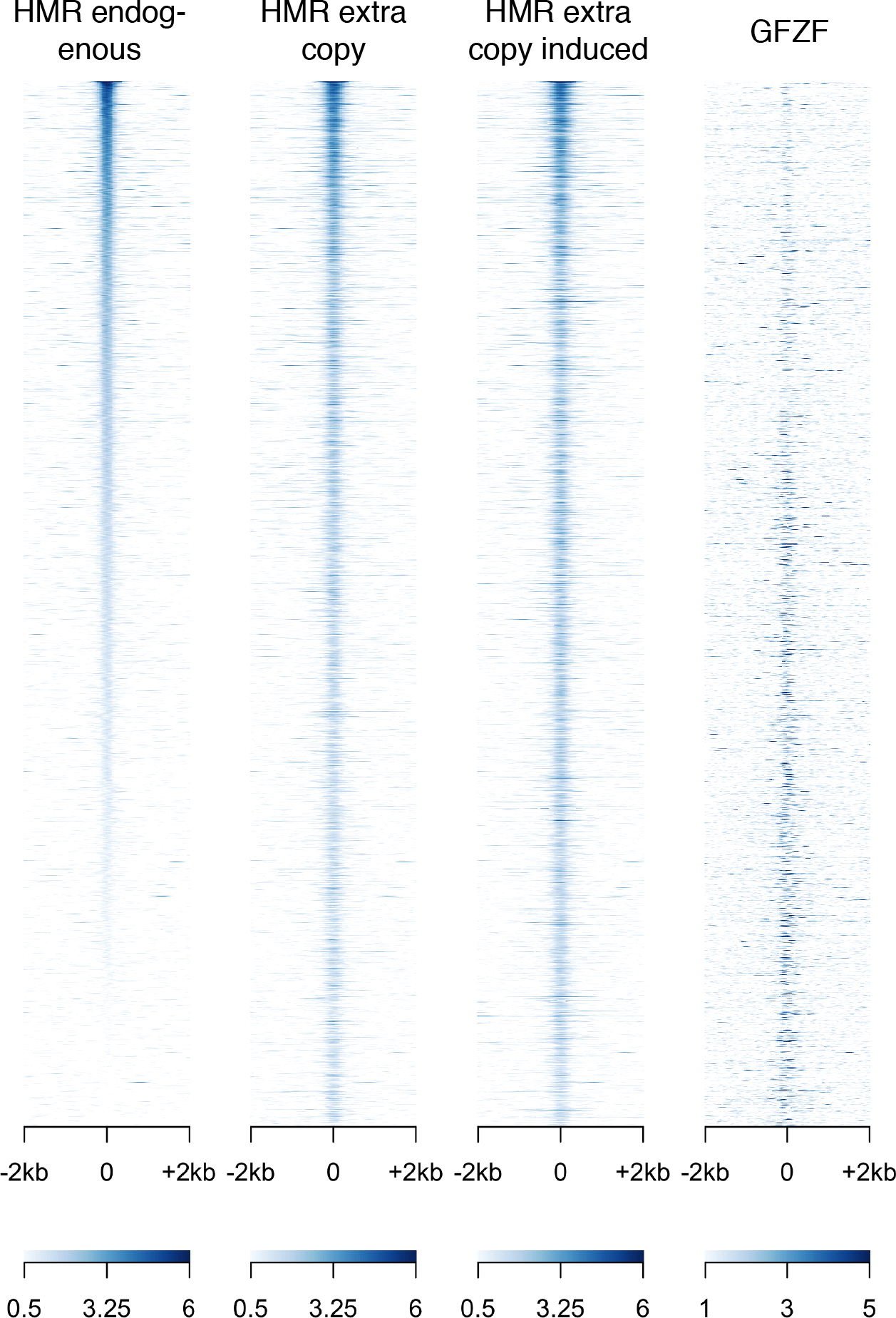
Heatmaps of HMR/GFZF signal all HMR bound sites in HMR over-expression experiments. The first three heat maps are for HMR signal. The final column represents the GFZF signal. All four columns are sorted by the strength of the HMR signal in the HMR over-expression induced state. When HMR is over-expressed, new sites appear near the end of the heatmap (as inferred by lack of HMR signal in the control).

**Supplemental Figure 9.**
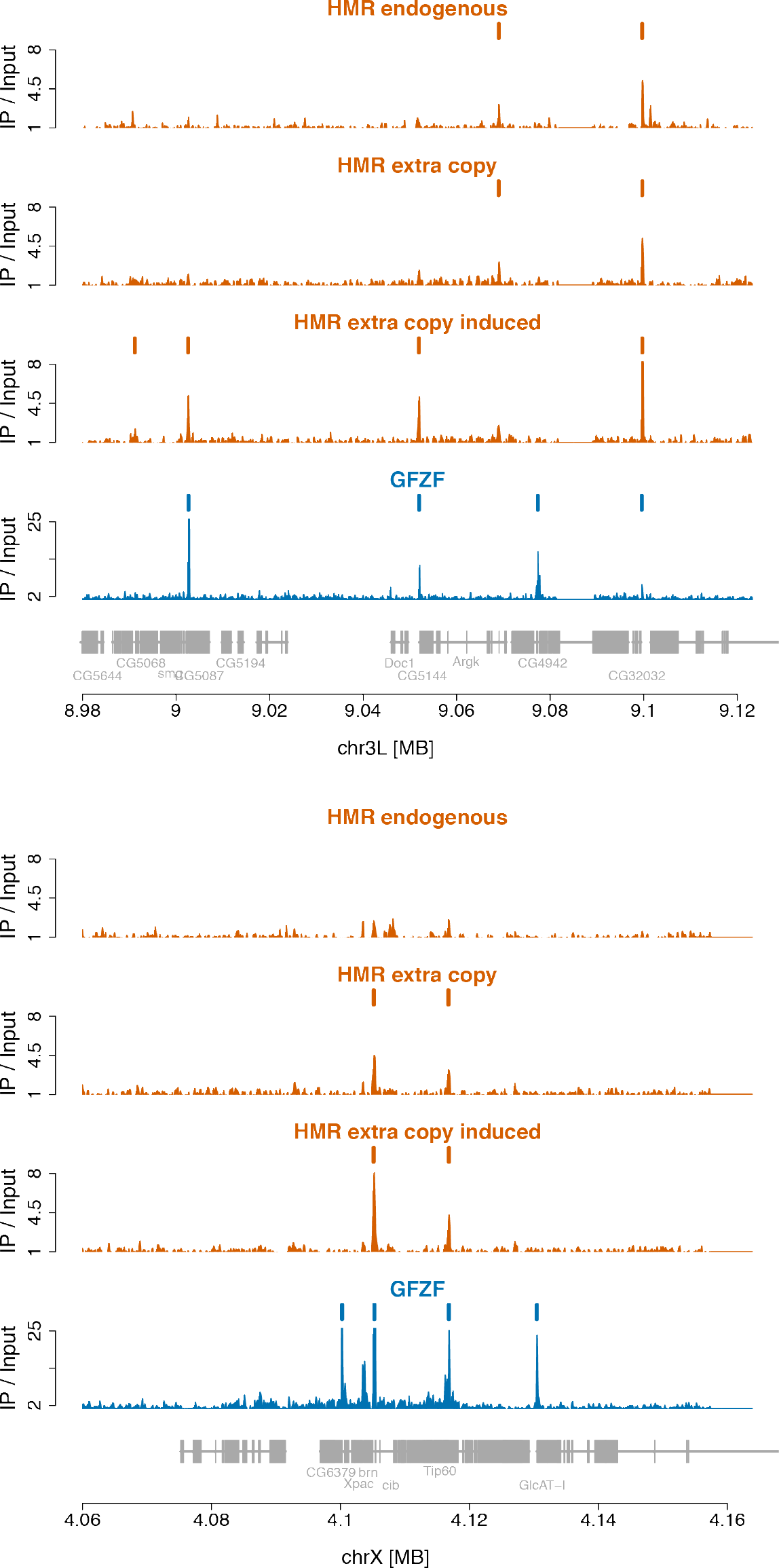
Additional Genome Browser windows for HMR over-expression. Additional windows to pair with Figure 5A. Peaks are marked with orange and blue bars for HMR and GFZF respectively.

**Supplemental Figure 10.**
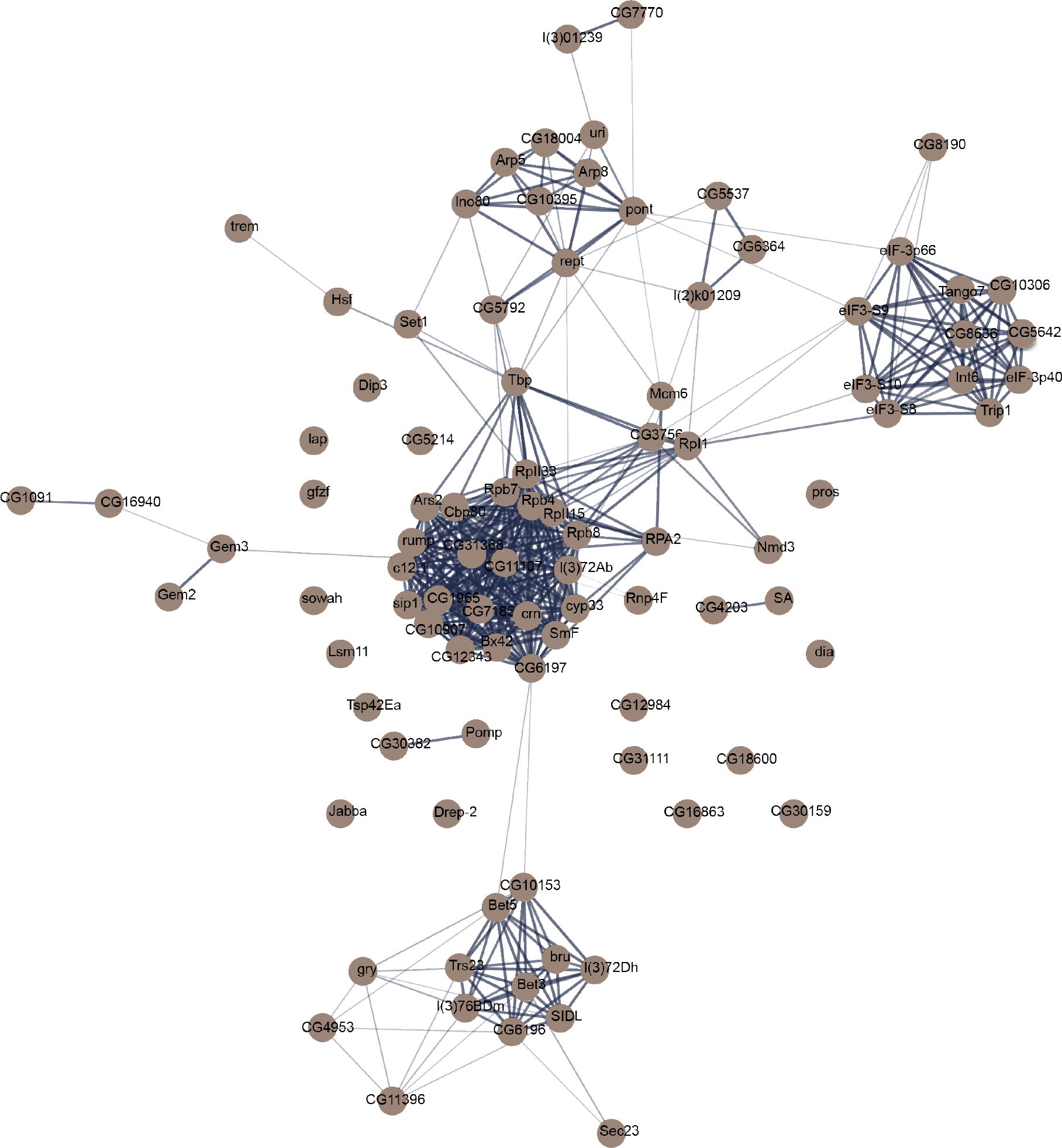
String analysis for the 96 genes identified in the GFZF proteome by AP-MS. Full analysis by STRING that was sampled from for Figure 6. Denser lines between genes represents a greater number of independent pieces of evidence that support the interaction.

